# CASBERT: BERT-Based Retrieval for Compositely Annotated Biosimulation Model Entities

**DOI:** 10.1101/2022.11.22.517475

**Authors:** Yuda Munarko, Anand Rampadarath, David P. Nickerson

**Affiliations:** Auckland Bioengineering Institute, University of Auckland, Auckland, New Zealand; The New Zealand Institute for Plant and Food Research Limited, Auckland, New Zealand

**Keywords:** BERT, Sentence-BERT, information retrieval, composite annotation embedding, RDF, ontology, BioModels, Physiome Model Repository (PMR)

## Abstract

Maximising FAIRness of biosimulation models requires a comprehensive description of model entities such as reactions, variables, and components. The COmputational Modeling in BIology NEtwork (COMBINE) community encourages the use of RDF with composite annotations that semantically involve ontologies to ensure completeness and accuracy. These annotations facilitate scientists to find models or detailed information to inform further reuse, such as model composition, reproduction, and curation. SPARQL has been recommended as a key standard to access semantic annotation with RDF, which helps get entities precisely. However, SPARQL is not suitable for most repository users who explore biosimulation models freely without adequate knowledge regarding ontologies, RDF structure, and SPARQL syntax. We propose here a text-based information retrieval approach, CASBERT, that is easy to use and can present candidates of relevant entities from models across a repository’s contents. CASBERT adapts Bidirectional Encoder Representations from Transformers (BERT), where each composite annotation about an entity is converted into an entity embedding for subsequent storage in a list-like structure. For entity lookup, a query is transformed to a query embedding and compared to the entity embeddings, and then the entities are displayed in order based on their similarity. The simple list-like structure makes it possible to implement CASBERT as an efficient search engine product, with inexpensive addition, modification, and insertion of entity embedding. To demonstrate and test CASBERT, we created a dataset for testing from the Physiome Model Repository and a static export of the BioModels database consisting of query-entities pairs. Measured using Mean Average Precision and Mean Reciprocal Rank, we found that our approach can perform better than the traditional bag-of-words method.

## 1 INTRODUCTION

In developing a biosimulation model, it is essential to provide a comprehensive description of the entities of the model related to processes, reactions, variables, mathematical equations and parameters. Formally, the COmputational Modeling in BIology NEtwork (COMBINE) community recommended the description in the form of semantic annotations using the Resource Description Framework (RDF) technology (Neal et al., 2019). With semantic annotations, complete and precise description is constructed in a composite manner involving various knowledge source terms and their relationships in a structured form (Gennari et al., 2011). Further, composite annotations are used as a community standard to encourage interoperability between platforms sharing and collaboration between modellers (Gennari et al., 2021; Welsh et al., 2021). This work supports this recommendation and goes beyond the composite annotation structure to provide an entity retrieval method with a simple data structure that is easy to deploy. Moreover, we suggest that the method is general enough so it can be implemented for different domains with composite annotations.

This complete and precise description is beneficial for understanding the model and subsequently becomes the key to rediscovery for verification and possible reuse. Verification that includes model curation ensures experimental results’ validity, reproducibility and consistency. Scientists can then confidently compare models or evaluate their proposed approaches. More comprehensive usability will enable model composition, creating larger-scale models, which is an essential aspect of modelling human physiology as a whole (Bassingthwaighte, 2000) and understanding the human body (Hunter et al., 2002).

The standard technology to locate and manipulate data stored in RDF format is SPARQL Protocol and RDF Query Language (SPARQL). SPARQL is powerful for retrieving data specifically and precisely (Pérez et al., 2009), although it requires a rigid syntax query. This rigidity becomes a barrier for most users even if they already have enough knowledge regarding the RDF triple and ontologies to explore RDF. For expert users, their queries still may fail caused by misspelling, capitalisation, and ontology terms variation. Therefore, text-based queries that can be composed freely as in commercial search engines are preferred, although the results are less precise.

The current state-of-the-art approaches offer a workaround by converting text-based queries to SPARQL by leveraging deep learning. Most of the works are for the question and answer purposes using generic graphs of RDF triples such as DBPedia ^1^, Yet Another Great Ontology (YAGO) ^2^, and Wikidata ^3^ and do not specifically support the searching of entities annotated compositely. Soru et al. (2020) and Yin et al. (2021) have considered the conversion as a language translation problem where SPARQL is the foreign language. Soru et al. (2020) implemented Long Short-Term Memory (LSTM) architecture to build sequence-to-sequence model and train the model over a dataset extracted from DBPedia. Then, Yin et al. (2021) extended the work by investigating the use of eight NMT methods built using Convolutional Neural Network (CNN), Recurrent Neural Network (RNN), and Transformer. CNN-based method (Gehring et al., 2017) performed the best within these methods, followed by Transformer-based (Vaswani et al., 2017). However, in the context of natural language translation, the use of Transformer is prospective to improve performance since it is not as mature as RNN and CNN. With the popularity of Bidirectional Encoder Representations from Transformers (BERT) (Devlin et al., 2018), Tran et al. (2021) created SPBERT, a Transformer-based model pre-trained using large DBPedia dataset for natural language to SPARQL and query results verbalisation tasks, and proved that the Transformer-based approach can surpass RNN and CNN. Adapting the created model for a new model is efficient by finetuning the existing model with less training data while keeping the property of the original model. Nevertheless, the text converted in these approaches must be in the natural language templates as formatted in the dataset. They cannot accommodate properly unstructured queries using keywords such as those used on commercial search engines.

Working well with structured and unstructured text-based queries, Natural Language Interface for Model Entity Discovery (NLIMED) provides an interface to retrieve entities annotated compositely (Munarko et al., 2022). It identifies phrases in the query associated with the physiological domain and links them to possible ontology classes and predicates. The link results then are composed as SPARQL and executed at the SPARQL endpoint to retrieve entities. A similar tool was developed by Sarwar et al. (2019), Model Annotation and Discovery (MAD), with the same domain but limited to entities in epithelial models. This limitation relates to template-based methods whose templates are customised for a particular topic, so the addition of topic coverage requires new templates.

This paper presents CASBERT, a method to retrieve entities in biosimulation models that are annotated compositely. We apply the information retrieval paradigm and leave the complexity of SPARQL providing a more expressive query composition while maximising the advantages of BERT. By adopting Sentence-BERT (Reimers and Gurevych, 2019), composite annotations describing entities are pre-calculated into entity embeddings and stored into a list-like structure. With the entity embeddings, a query that is also converted into a query embedding is compared using a similarity formula; then, the ranked results are displayed.

Recently, the biosimulation model formats in the two largest repositories, the Physiome Model Repository (PMR) (Yu et al., 2011) and BioModels Database (Chelliah et al., 2015), have used RDF to describe their entities. Composite annotations are largely used to describe models in CellML (Cuellar et al., 2003) and SBML (Hucka et al., 2003) formats, detailing entities with terms in ontologies such as anatomical location, chemical compound, physics of biology, gene, and protein. We generated test data from these repositories in the form of query and entities pairs. CASBERT performance was measured using Mean Average Precision and Mean Reciprocal Rank, and compared to the traditional bag-of-words method, the score was significantly higher. In addition, a list-like structure for storing embedding entities provides the flexibility to manage the addition, subtraction, or insertion of entities so that implementation as a search engine is possible. CASBERT can also be implemented for composite annotation search in various domains such as chemistry, pharmacy, and medical. Our implementation, dataset, and experiment setup are publicly available ^4^.

## 2 MATERIALS AND METHODS

CASBERT provides approaches to convert composite annotations used to define entities and queries to embeddings. Entity embeddings are pre-calculated and stored in a list-like structure, whereas a query embedding is created on the fly when a request is made. Entities then are presented in order from the most relevant by initially calculating the similarity values between query embedding and entity embeddings.

The conversion to embedding methods are based on Bidirectional Encoder Representations from Transformers (BERT) (Devlin et al., 2018) by implementing Sentence-BERT (Reimers and Gurevych, 2019). For the experiment, we created a dataset by collecting composite annotations from biosimulation models from the PMR and BioModels database. These conversion methods are described in the following subsections, starting with the dataset used.

### 2.1 Biosimulation Model -Composite Annotation Query (BM-CAQ) Dataset

We constructed the dataset for the experiment by extracting compositely annotated entities in the PMR and BioModels database. From the RDF and CellML files in the PMR, we got 4,652 entities with textual descriptions. These textual descriptions are usually short, concise, and created by experts in biosimulation modelling; therefore, these descriptions can represent the actual queries used to find the entities. Altogether, 338 unique textual descriptions were considered queries related to at least one entity. Moreover, we extended the number of queries by randomly inserting terms in predicates to create a new query, so we have 534 additional queries. The first query set is called *noPredicate*, and the latter is called *withPredicate*. We applied the same strategy to extract queries from BioModels database, which returned 834 *noPredicate* and 1,541 *withPredicate* queries. In its entirety, this dataset is named Biosimulation Model -Composite Annotation Query (BM-CAQ) and sample data are shown in Supplementary Material, see Tables S1 to S4.

### 2.2 Composite Annotation Search Using BERT (CASBERT)

We used BERT (Devlin et al., 2018) and Sentence-BERT (Reimers and Gurevych, 2019) to convert queries and entities to embeddings and to classify the query to be compared to the appropriate query embedding list. BERT provides pre-training models that are built based on massive corpora that can be fine-tuned with a smaller corpus to maximise performance for domain-specific uses. A sentence is converted into embedding by splitting to tokens, then calculating the embedding of each token unique to the context, such as the surrounding tokens and position. Therefore, the same token in different sentences will have a different embedding. This attention to context correlates with high reliability in several natural language processing tasks, such as named entity recognition, concept extraction, and sentiment analysis, and is relatively better than non-context embedding (Taillé et al., 2020; Arora et al., 2020). In addition, the form of the token, which is part of the sentence, makes BERT more adaptive to typographical errors and variations of word writing.

The most straightforward approach to create a sentence embedding is by averaging the token embeddings, however this often leads to poor performance (Reimers and Gurevych, 2019). Resolving this issue, Sentence-BERT offers a better concatenation method optimised for Semantic Textual Similarity (STS) by applying Siamese (Bromley et al., 1993) and Triplet loss (Weinberger and Saul, 2009) networks.

CASBERT converts the entity’s composite annotation and query to embeddings by adopting Sentence-BERT to transform each ontology class or predicate concept in the composite annotation or the query to an embedding. These concept embeddings then are combined, producing entity embedding or query embedding. However, establishing entity embedding and query embedding is different, explained as follows.

#### 2.2.1 Entity Embedding

Here we use an entity instance of the *GAP*_*g*_ variable in the brain energy metabolism model (Cloutier et al., 2009) for which the CellML model is available in the PMR^5^. Figure 1A shows the composite annotation regarding the concentration (OPB:00340) of glyceraldehyde 3-phosphate (CHEBI:17138) in an Astrocyte (FMA:54537). There are five triples with the subject and objects: root (*GAP*_*g*_*/GAP*_*g*_), ontology classes (OPB:00340, CHEBI:17138, FMA:54537), and intermediate subjects/objects (entity 1, entity 2). Figure 1B presents interconnected triples creating paths that display a clear relationship between root and ontology classes. The terms used to refer as intermediate subject/object are usually generic and are similar across all composite annotations, so they cannot be used as a differentiator; therefore, we ignore them (Figure 1C). Next, we remove predicates that directly connect intermediate subjects, e.g. ‘is’, and ontology classes, because these only describe the intermediate subjects/objects, not the entity.

**Figure 1.**
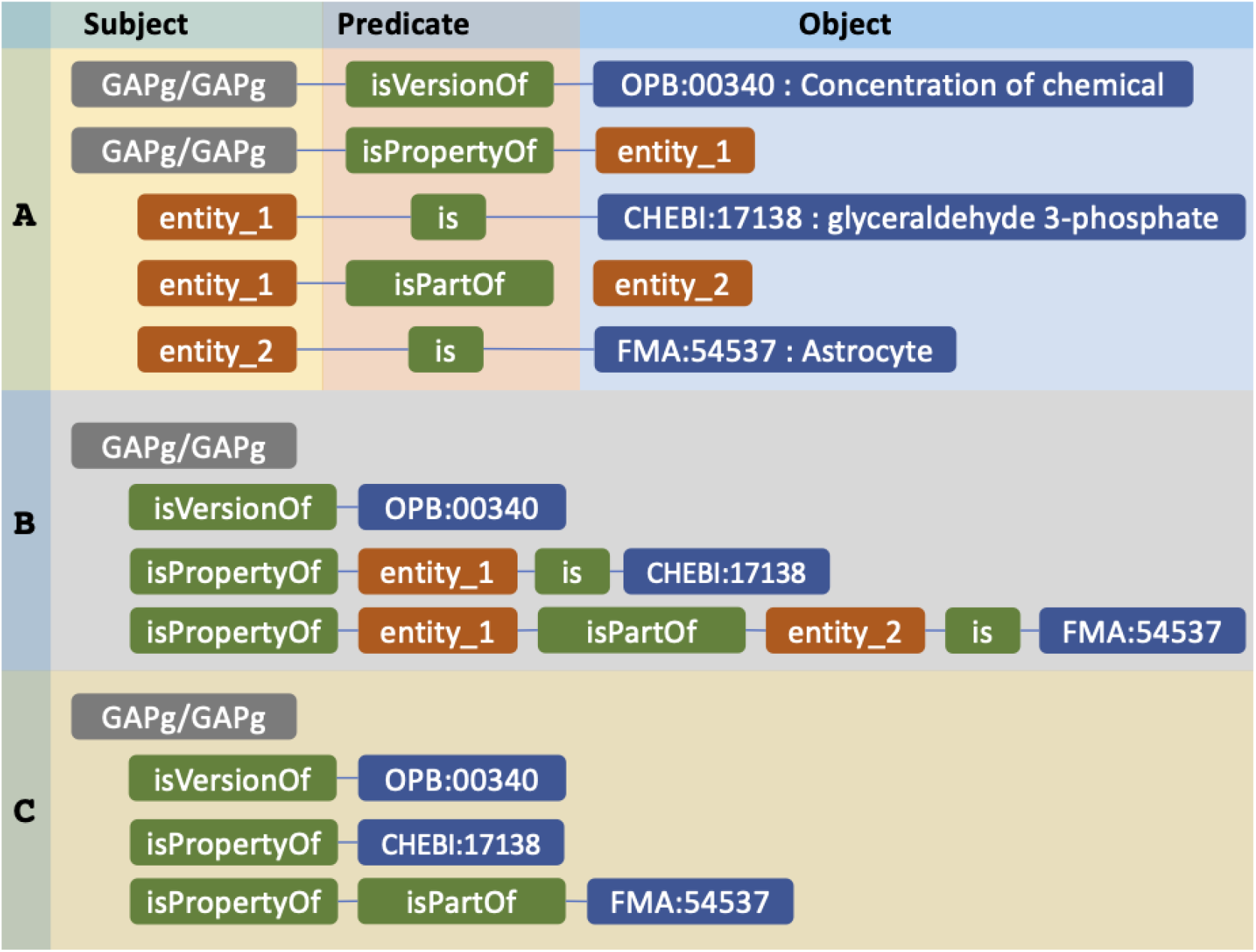
The example of an entity compositely annotated using RDF and its representation for further conversion to embedding. (A) The composite annotation of *GAP*_*g*_*/GAP*_*g*_ entity by three ontology classes. (B) Paths consist of predicates connecting the entity to ontology classes. (C) The representation of the entity before converted to entity embedding.

Figure 2 illustrates the translation of a composite annotation to embedding. Initially, CASBERT calculates the embedding of each path *e*_*pt*_ by combining its ontology class embedding *e*_*c*_ and the average of predicate embeddings *e*_*p*_ using Equation (2). *e*_*c*_ is the average of ontology class feature embeddings, including preferred label embedding and synonym embedding (see Equation (1)). We do not use other features such as parent label and definition because the use of the preferred label and synonym only can give a higher performance (Munarko et al., 2022). We apply *w*_*p*_ between 0 to 1 in Equation 2 as a multiplier to limit the role of *e*_*p*_ to the path embedding, so it does not exceed the ontology class embedding. Finally, all path embeddings are averaged to get entity embedding *e*_*e*_ as presented by Equation 3. Furthermore, all annotated entities in the PMR are converted into embeddings and arranged in a list-like structure to facilitate create, read, update, and delete (CRUD) operations.

**Figure 2.**
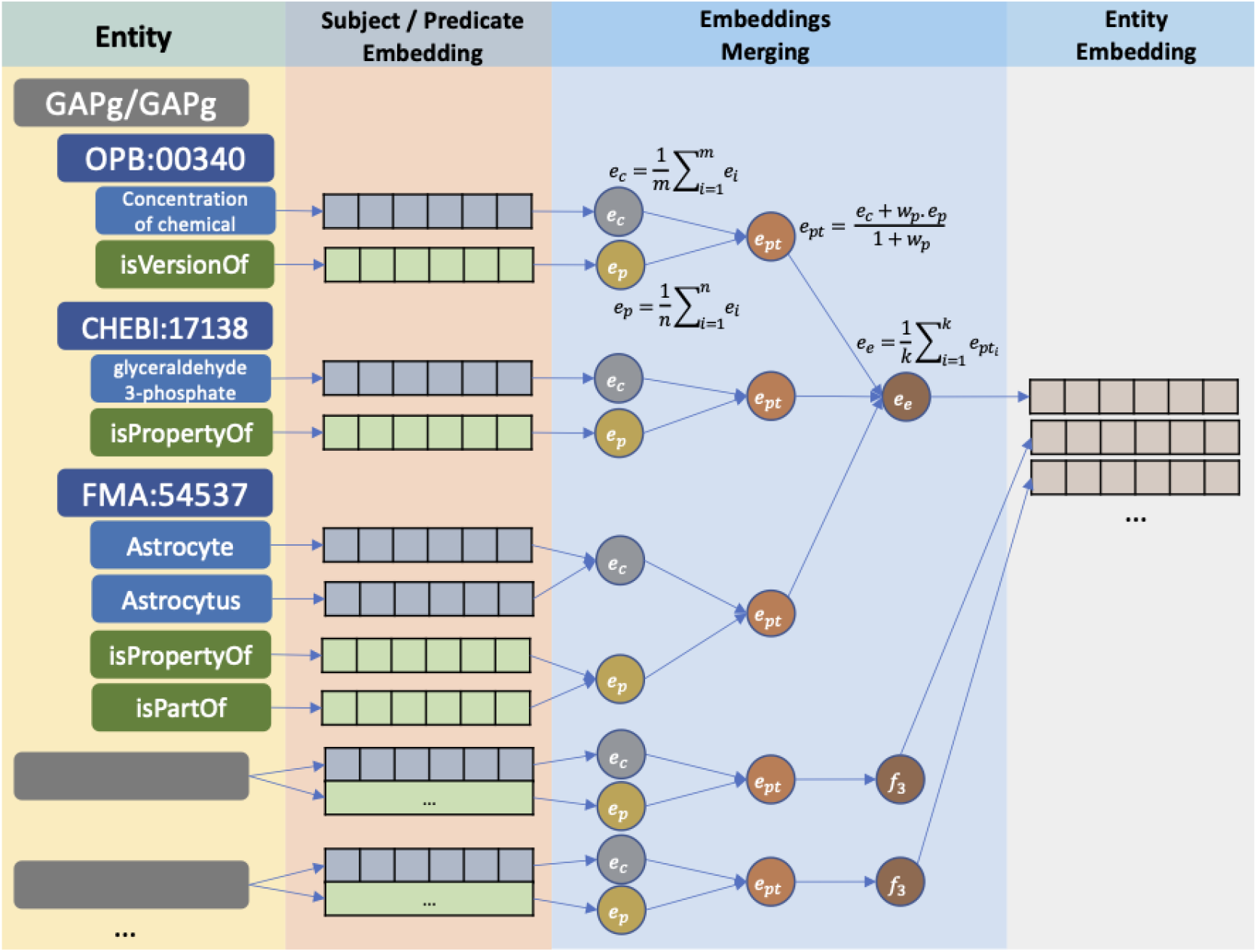
The conversion of entities to entity embeddings. For each entity, ontology classes and predicates are encoded into subject and predicate embeddings and then combined into one-dimensional embeddings.

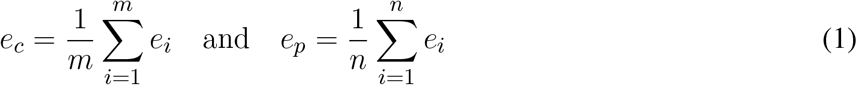

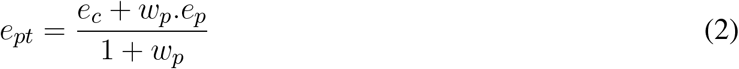

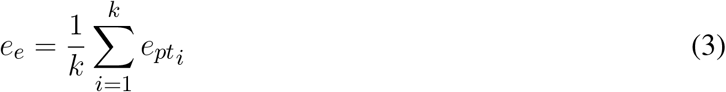

Ontology class feature embedding and predicate embedding are generated using Sentence-BERT. While converting the ontology class features is straightforward, the predicate is not. The predicate should be preprocessed by removing the leading substring leaving the primary term such as ‘isVersionOf’ and ‘isPartOf. Further, this primary term is normalised into a phrase such as ‘is version of’ and ‘is part of’ and finally converted into embedding.

#### 2.2.2 Query Embedding

To search for the *GAP*_*g*_ variable in Figures 1, let’s say the user queries ‘triose phosphate concentration in astrocytes’. A query can be thought of as a composite annotation summary involving concepts related to ontology classes and connecting predicates. Therefore, as presented in Figure 3, the ontology class concepts can be identified as ‘concentration’, ‘triose phosphate’, and ‘astrocyte’, where ‘triose phosphate’ is a synonym for ‘glyceraldehyde 3-phosphate’. CASBERT implements SciSpacy (Neumann et al., 2019) for the identification with the ‘en core sci scibert’ as Named Entity Recognition (NER) model suitable for biomedical data. These ontology class concepts then are converted to embeddings and combined by averaging (Equation (4)). Since most of the concepts are loosely related to the ontology class with a similarity value of less than 1, the combined embedding is normalised by *w*_*ph*_ which is the average of the maximum similarity of each concept *p* to ontology classes *ocs*. This similarity calculation follows Equation (6).

**Figure 3.**
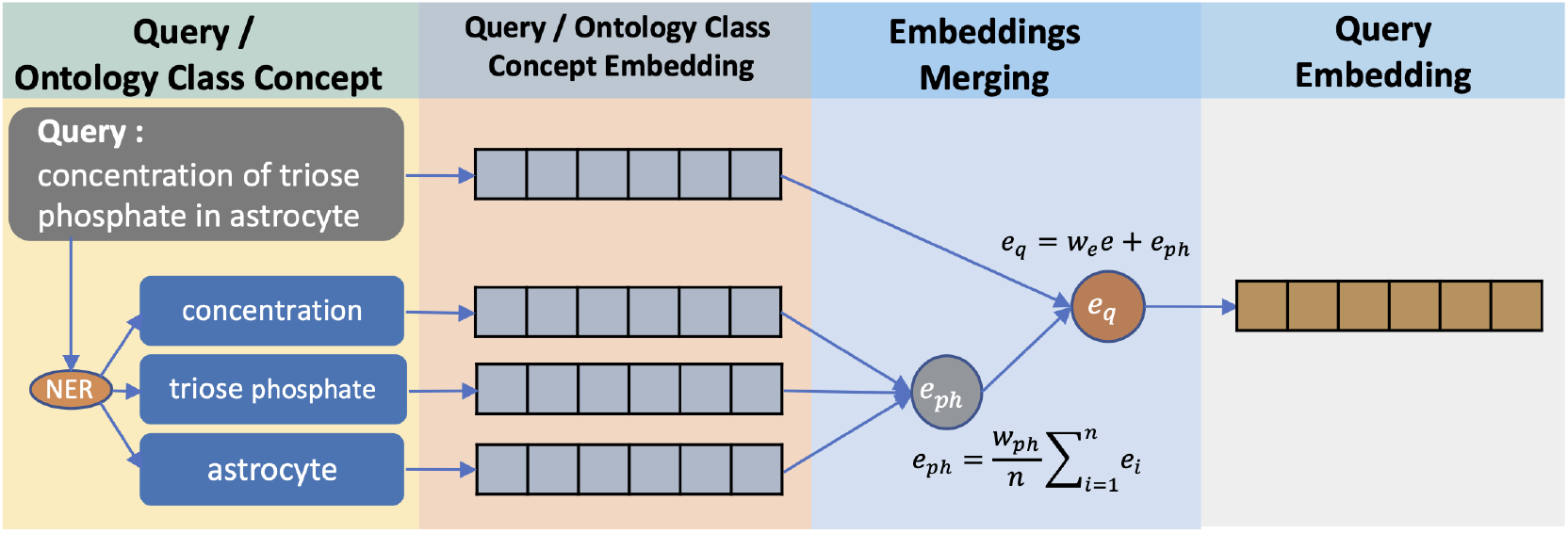
The conversion of a query to query embedding. The query is chunked into phrases related to the ontology class concept and then converted into embeddings using Sentence-BERT. These phrase embeddings are then averaged and combined with the entire query text embedding.

Unlike the ontology class concepts, the predicate concepts are not always explicitly represented using phrases; instead, they are usually implicitly encoded and represented using general conjunctions and prepositions. Therefore, to capture ontology classes and their relationship patterns via predicates, we also add an embedding of whole query terms. Hence, the query embedding is obtained by converting the ontology class concept phrases and the entire query terms to embeddings, *e*_*ph*_ and *e* consecutively, where *e* is multiplied with the empirically decided weight *w*_*e*_, and then combining them using the addition function, Equation (5).

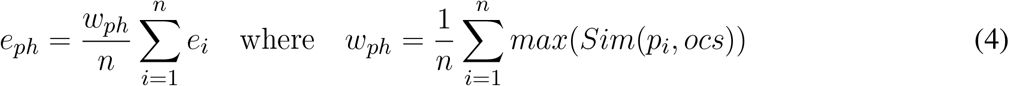

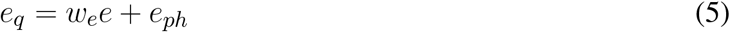

For experimental purposes, we augment the *e*_*p*_*h* calculation with the predicate phrases identified using a similar method implemented in NLIMED (Munarko et al., 2022). These phrases are converted into embeddings and averaged based on the related ontology class. Then the ontology class embedding and the predicate embedding mean are combined where the latter is multiplied by *w*_*p*_ as in Equation 2.

#### 2.2.3 Entity Retrieval

With the availability of the entity embedding list and query embedding, we can now calculate their similarities and present the relevant entities sorted from the highest value as illustrated in Figure 4.

**Figure 4.**
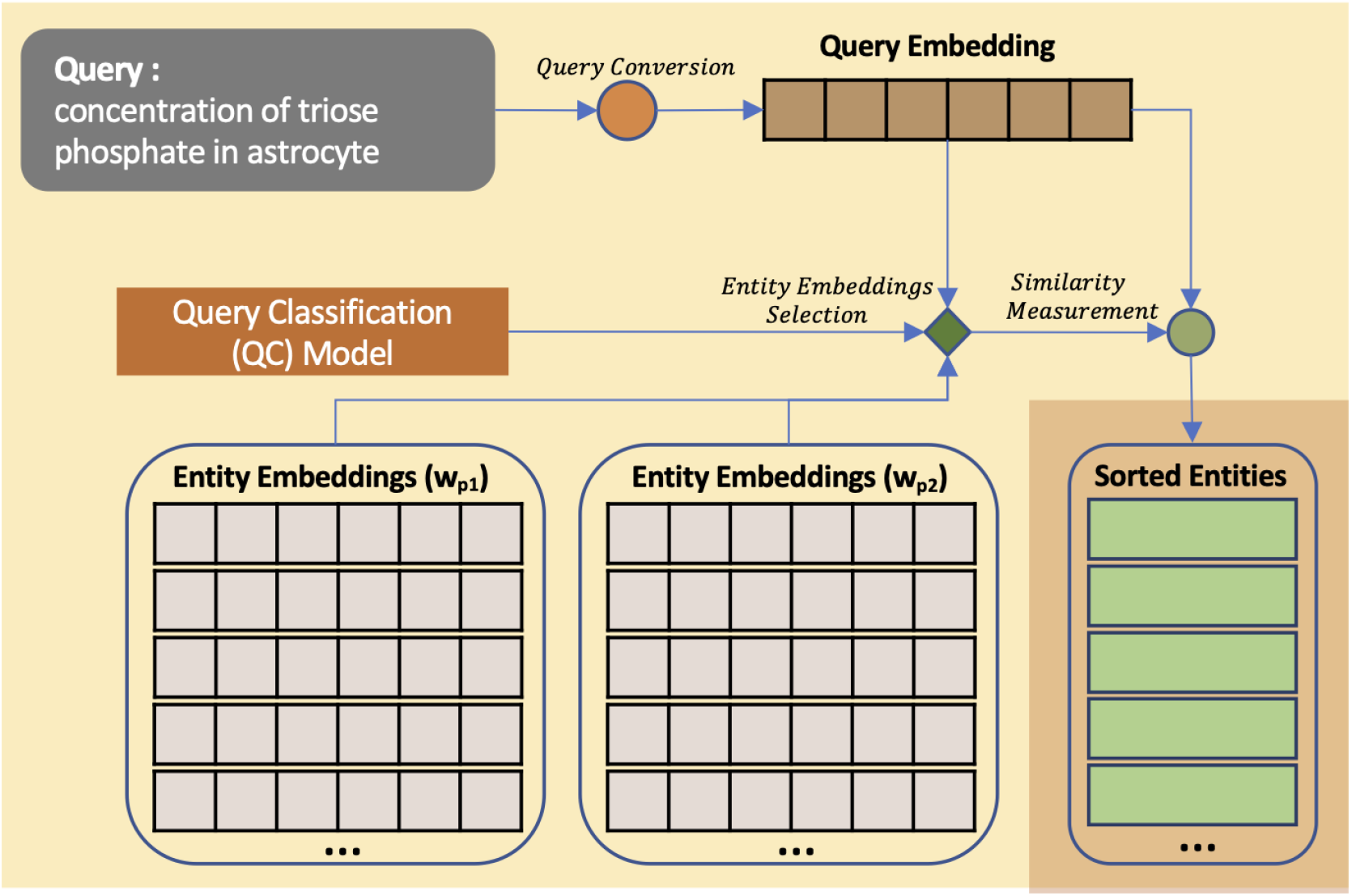
Query to entities matching using CASBERT. After converting the query to a query embedding, the entity embedding list to be compared with the query embedding is selected using Query Classification Model. Relevant entities then are retrieved using cosine similarity measure.

##### Query - Entity Similarity

We implemented Cosine Similarity (CS) (Salton and McGill, 1983) to calculate the similarity value between a query embedding *e*_*q*_ and an entity embedding *e*_*e*_. CS of two embeddings is the dot product of both embeddings divided by the multiplication of the magnitude of both embeddings (Equation 6). Thus, CS ignores the magnitude of each embedding, making it suitable for the high dimensionality nature of embedding. Additionally, ‘multi-qa-MiniLM-L6-cos-v1’ pre-trained sentence transformer model (Reimers and Gurevych, 2019) used to convert a sentence to an embedding in CASBERT is optimised with CS.

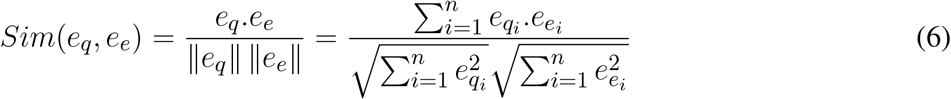

##### Query Classification

Considering that we can create multiple embedding lists of entities with different *w*_*p*_ (see Equation 2), we found that some queries can retrieve best when compared to a list with *w*_*p*_ = 0, while others to a list with *w*_*p*_ *>* 0. Therefore, we created the Query Classification (QC) model to select the appropriate list to calculate query-entities similarity. As an experiment, we created two lists with *w*_*p*_ = 0 and *w*_*p*_ = 0.22. The QC model was trained using Transformers (Wolf et al., 2020) and the ‘bert-base-uncased’ pre-trained model where the training data was initially augmented using nlpaug ^6^ to increase its size and diversity. The training process in more detail is presented in Supplementary Figure S1.

## 3 EXPERIMENTS AND RESULTS

### 3.1 Experiment Setup

We conducted experiments to understand the behaviour of CASBERT implemented using different retrieval strategies against different query types. We believe that using all terms in the query alone without paying attention to phrases related to the concept of ontology classes will be more efficient in execution time; however, combining the two is likely to improve retrieval quality.

The experiment used our BM-CAQ dataset including *noPredicate* and *withPredicate* query sets and the combination of both. We varied the retrieval method by combining terms in the query as a whole and phrases related to the concept of ontology classes. We created two entity embedding lists with *w*_*p*_ = 0 and *w*_*p*_ = 0.22 where 0.22 is chosen randomly to indicate the role of the predicate in different retrieval methods. Then we also compared the retrieval methods that implement the query classifier and those that do not. The query embedding is created according to Equation (5) using *w*_*e*_ = 1.9. We also measured the performance of BM25 (Robertson and Walker, 1994), a bag-of-words method, as the gold standard. The description of the retrieval method and query set in this experiment is presented in Table 1.

**Table 1.**
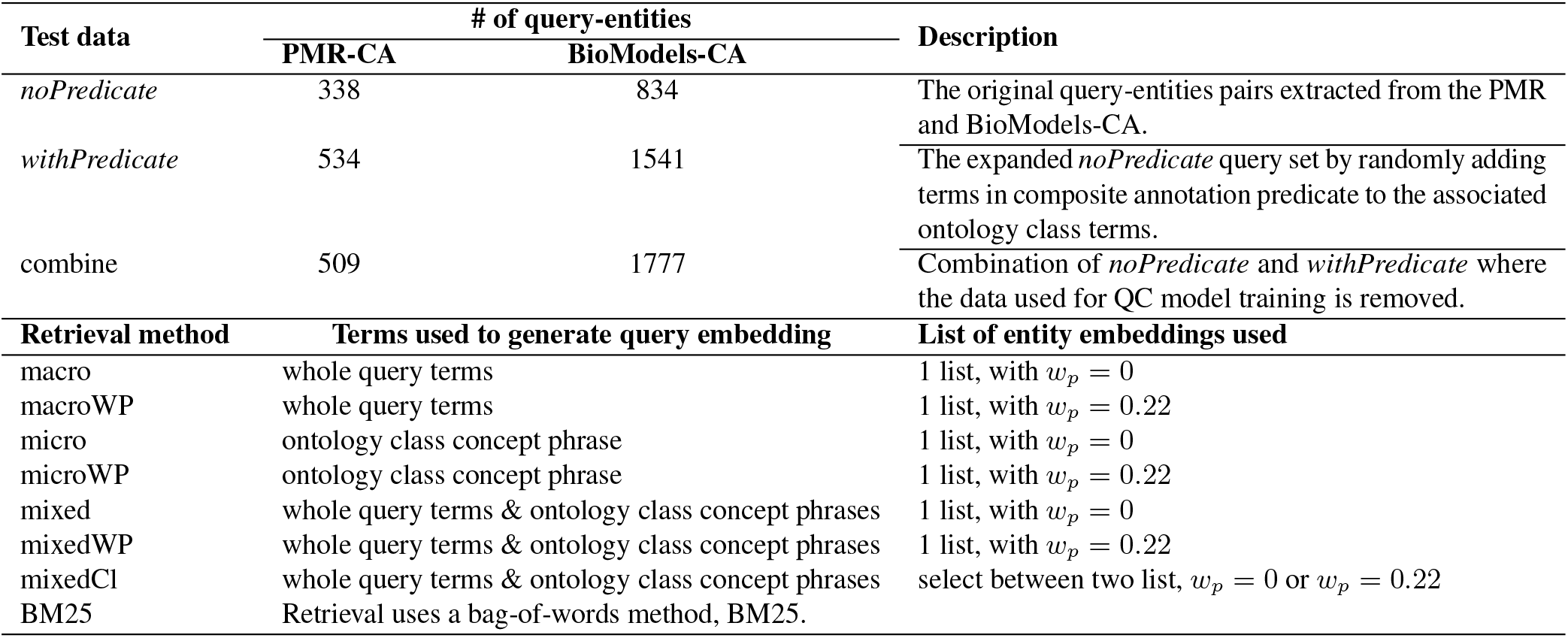
The strategies to measure CASBERT performance. There are three query sets and eight retrieval methods including BM25 as the gold standard.

| Test data | # of query-entities |  | Description |
| --- | --- | --- | --- |
|  | PMR-CA | BioModels-CA |  |
| *noPredicate* | 338 | 834 | The original query-entities pairs extracted from the PMR and BioModels-CA. |
| *withPredicate* | 534 | 1541 | The expanded *noPredicate* query set by randomly adding terms in composite annotation predicate to the associated ontology class terms. |
| combine | 509 | 1777 | Combination of *noPredicate* and *withPredicate* where the data used for QC model training is removed. |
| Retrieval method | Terms used to generate query embedding |  | List of entity embeddings used |
| macro | whole query terms | | 1 list, with $w_p = 0$ |
| macroWP | whole query terms | | 1 list, with $w_p = 0.22$ |
| micro | ontology class concept phrase | | 1 list, with $w_p = 0$ |
| microWP | ontology class concept phrase | | 1 list, with $w_p = 0.22$ |
| mixed | whole query terms & ontology class concept phrases | | 1 list, with $w_p = 0$ |
| mixedWP | whole query terms & ontology class concept phrases | | 1 list, with $w_p = 0.22$ |
| mixedCI | whole query terms & ontology class concept phrases | | select between two list, $w_p = 0$ or $w_p = 0.22$ |
| BM25 | Retrieval uses a bag-of-words method, BM25. |  |  |

### 3.2 Evaluation Metric

We measured CASBERT performance for each query set *Q* using Mean Average Precision for the top k results (*mAP* @*k*), Equation 8. *mAP* @*k* is based on Average Precision at k (*AP* @*k*) as shown by Equation 7, where *R* is the number of relevant entities in the results, *P* @*i* is the proportion of relevant entities in the top *i* results, and *r*@*i* is a relevance function that returns 0 or 1 for the irrelevance or relevance of the entity at position *i*, respectively. Furthermore, we set the value of *k* to 10 because search results are usually arranged in pages of 10 entities, and users are generally only interested in the first page.

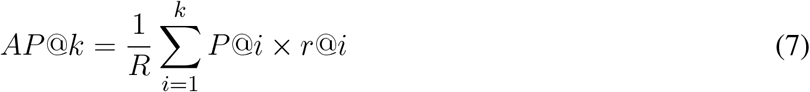

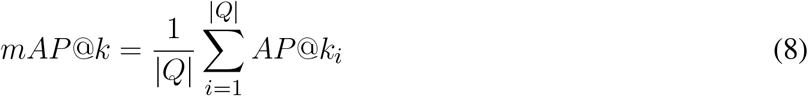

Moreover, we also use Mean Reciprocal Rank (*mRR*) with Equation 9 measuring the mean of the multiplicative inverse of the first entity in the results found to be relevant (*ranks*_*i*_). This measure is also helpful since some of the queries in BM-CAQ are associated with one entity only.

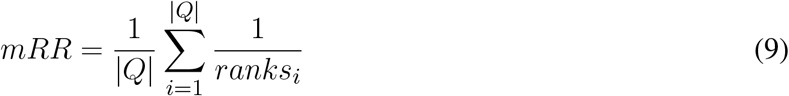

### 3.3 Results

From Table 2, we can see that all of the methods used in CASBERT have higher *mAP* @10 and *mRR* than the gold standard BM25. Strategies that combine embeddings of the whole query terms and phrases related to the concept of ontology classes (mixed, mixedWP, mixedCl) perform best compared to the others (indicated by bold values). Moreover, in the mixedCl, the use of the QC model to select an entity embedding list can slightly improve the retrieval quality and make it the best method. This better performance confirms our initial presumption, although the use of whole query terms only can retrieve entities appropriately.

**Table 2.** CASBERT performance over three test data types and seven searching strategies compared to the bag-of-words method (BM25) measured using *mAP* @10 and *mRR*. The numbers of entities in the PMR and BioModels are 4,652 and 54,456 respectively.

| Method | *noPredicate* |  | *withPredicate* |  | *combine* |  |
| --- | --- | --- | --- | --- | --- | --- |
| | $mAP@10$ | $mRR$ | $mAP@10$ | $mRR$ | $mAP@10$ | $mRR$ |
| **PMR-CA** |  |  |  |  |  |  |
| macro | 0.645216 | 0.643424 | 0.572291 | 0.597649 | 0.601406 | 0.602604 |
| macroWP | 0.626565 | 0.624992 | 0.585014 | 0.606947 | 0.604012 | 0.608011 |
| micro | 0.619483 | 0.617890 | 0.534427 | 0.549304 | 0.592228 | 0.595516 |
| microWP | 0.606255 | 0.608560 | 0.504055 | 0.515426 | 0.574869 | 0.579472 |
| mixed | 0.656636 | 0.656326 | 0.572109 | 0.583003 | 0.623287 | 0.624638 |
| mixedWP | 0.641843 | 0.641940 | 0.585494 | 0.612151 | 0.615885 | 0.621419 |
| mixedCI | **0.660346** | **0.659399** | **0.589960** | **0.615694** | **0.626420** | **0.631739** |
| BM25 | 0.459375 | 0.443034 | 0.464496 | 0.487373 | 0.456505 | 0.457420 |
| **BioModels-CA** |  |  |  |  |  |  |
| macro | 0.345486 | 0.340106 | 0.310285 | 0.302129 | 0.356034 | 0.347905 |
| macroWP | 0.330100 | 0.326029 | 0.326831 | 0.325134 | 0.365903 | 0.362892 |
| micro | 0.294353 | 0.291914 | 0.283026 | 0.283291 | 0.312949 | 0.309471 |
| microWP | 0.301564 | 0.296179 | 0.274491 | 0.271451 | 0.314506 | 0.309211 |
| mixed | 0.347681 | 0.345045 | 0.326664 | 0.323393 | 0.370148 | 0.365433 |
| mixedWP | 0.335051 | 0.331222 | **0.344296** | **0.344932** | 0.379058 | **0.377573** |
| mixedCI | **0.348054** | **0.344794** | 0.343567 | 0.344058 | **0.379351** | 0.377557 |
| BM25 | 0.240671 | 0.231515 | 0.294758 | 0.294779 | 0.280370 | 0.277879 |

Illustrating the effect of different *mAP* @10 and *mRR* values, Figure 5 show the top 10 retrievals of entities for three query examples retrieved using mixedCl and BM25. For the first and the third examples, mixedCl and BM25 retrieved the same number of relevant entities; however, the relevant entities appear in higher ranks with mixedCl rather than BM25. For the second example, mixedCl can retrieve more relevant entities and rank better. Hence, higher *mAP* @10 and *mRR* provide a better search experience with relevant entities being ranked higher or more relevant entities being presented.

**Figure 5.**
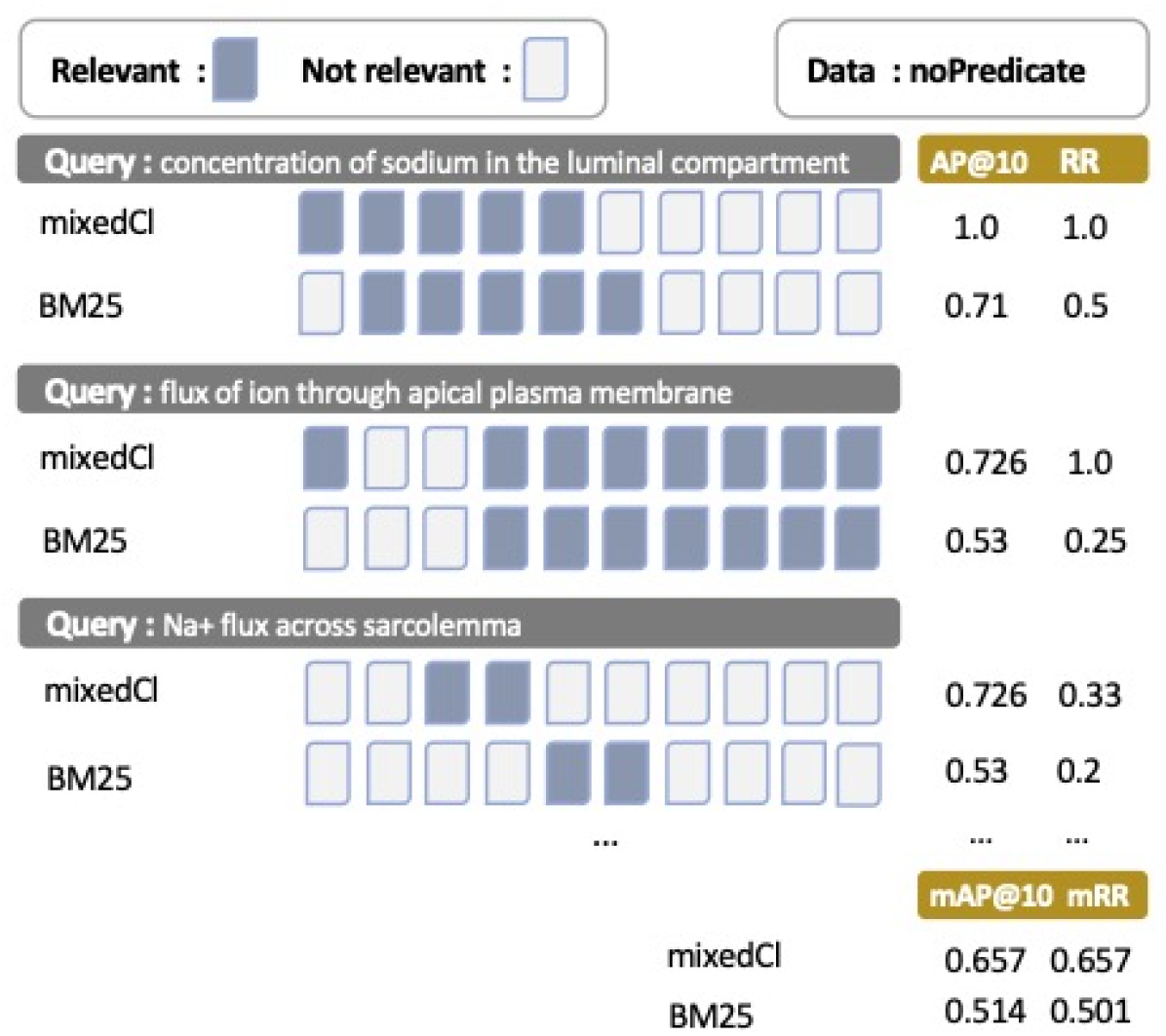
The illustration of how query results are rendered, where left side is more relevant than right side. This figure presents query-results examples from *noPredicate* test data using mixedCl and BM25 with *AP* @10, *RR, mAP* @10, and *mRR* values.

## 4 DISCUSSION

We have demonstrated that the adaptation of BERT based embedding to encode an entity’s composite annotation and query in CASBERT can surpass the performance of the standard bag-of-words methods. This better performance is closely related to the way BERT tokenises an input sentence and converts the token into embedding. BERT implements WordPiece tokeniser (Wu et al., 2016) which properly accommodates variation in wording, spelling error, and white space missing issues. Therefore, unique words that appear in specific domains such as biology can be segmented into appropriate tokens. Furthermore, BERT’s embedding process considers contexts, including the surrounding tokens and the token’s position, so that the same token can have a different embedding in different sentences. Hence, the combination of all token embeddings can appropriately represent a sentence. Moreover, contexts contribute to the generation of similar embeddings for tokens with synonymy meaning.

In the following subsections, we discuss CASBERT improvements, analysed based on different embedding generation methods. Then the discussion is followed by the possibility of implementing CASBERT in various domains and complementing SPARQL. Finally, we present our recommendations and future works.

### 4.1 Performance Increase

As presented in Table 2, our proposed methods are quite effective in converting entity’s composite annotation and query to embeddings and retrieving relevant entities based on the provided query. The analysis of those methods and performance based on similarity value between query and composite annotation is described below.

#### 4.1.1 Entity’s Composite Annotation to Embedding Methods

We have experimented with creating entity embedding by considering predicates (*w*_*p*_ *>* 0) and not (*w*_*p*_ = 0). Entity embeddings considering predicates are mostly suitable for queries containing the term predicate or its synonyms, though not all. In contrast, the ones without predicates usually are more suitable for queries without predicate terms. In contrast, the ones without predicates usually are more suitable for queries without predicate terms. Intuitively, identifying the availability of predicate related terms in the query and then selecting the correct entity embedding based on this availability is possible, however, this method does not perform as expected; therefore, we created a Query Classification (QC) model to select entity embedding based on the query. Applying the QC model with the mixed method, retrieval performance increases quite convincingly with three lower *mAP* @10 and *mRR* measurements but nine higher for the other measurements (Table 2, bold values). While only two *w*_*p*_ values, 0 and 0.22, are used in this experiment, we predict that *w*_*p*_ should be adaptive to the query, so in the future, we recommend specifying *w*_*p*_ automatically. However, this adaptive *w*_*p*_ approach will sacrifice the simplicity of the current entity’s composite annotation list because ontology class and predicate embeddings should be separately managed, and there should be a mechanism to create entity embedding with given *w*_*p*_ value effectively.

#### 4.1.2 Query to Embedding Methods

The empirical results show that the mixed method performs best, followed by macro and micro methods consecutively. The micro method is intended to detect ontology class concepts in a query and generate a query embedding by combining all concept embeddings. However, the detection accuracy depends on the NER method’s performance in identifying the concept of ontology classes and the query created by the user.

The macro method is convincingly better than the micro method. This higher performance might be related to encoding whole query terms containing all ontology class concepts, including their relationship in a single embedding unit. However, individual ontology class concepts are not considered, allowing slight entity detection inaccuracies. Overall, this method is the most efficient because it only performs a one-step conversion from query to embedding, in contrast with the other methods that identify multiple ontology class concepts and convert to embeddings and then combine them.

The mixed-method can slightly increase *mAP* @10 and *mRR*. As expected, this merge takes good account of the macro method’s advantages and emphasises the critical ontology class concepts provided in the query. To avoid the decisive role of the ontology class concepts, delimiting with *w*_*ph*_ (Equation (4)) can give adequate proportion. Although this method is not the most efficient, its computational cost is linearly increased depending on the identified ontology class concepts. Due to its highest effectiveness, we recommend the mixed-method to be implemented for composite annotation search.

#### 4.1.3 Performance Analysis Based on The Similarity of Query and Entity’s Composite Annotation

Figure 6 shows CASBERT’s ability in retrieving entities for various queries differentiated by their similarity to relevant entities for PMR-CA. We calculated the similarity directly using the query embedding generated with the macro method against the entity embedding. CASBERT performance is higher than BM25 when the similarity value is more than 0.3 and achieves the highest margin for similarity from 0.5 to 0.9, covering about 96% of the total test data. This pattern indicates CASBERT benefits because most user queries fall within this range. For very low query-entity similarity, 0.1 to 0.2, BM25 is better because a limited number of the same terms, one or two, can direct the query to relevant entities. In contrast, embedding in CASBERT may lead the query to entities with different terms but having the same context. Unfortunately, the combination of a low similarity value and the absence of a common term results in lower performance. Furthermore, we found a similar pattern for BioModels-CA, although we do not measure performance for low query-entity similarity values (see Supplementary Figure S2 and Table S5).

**Figure 6.**
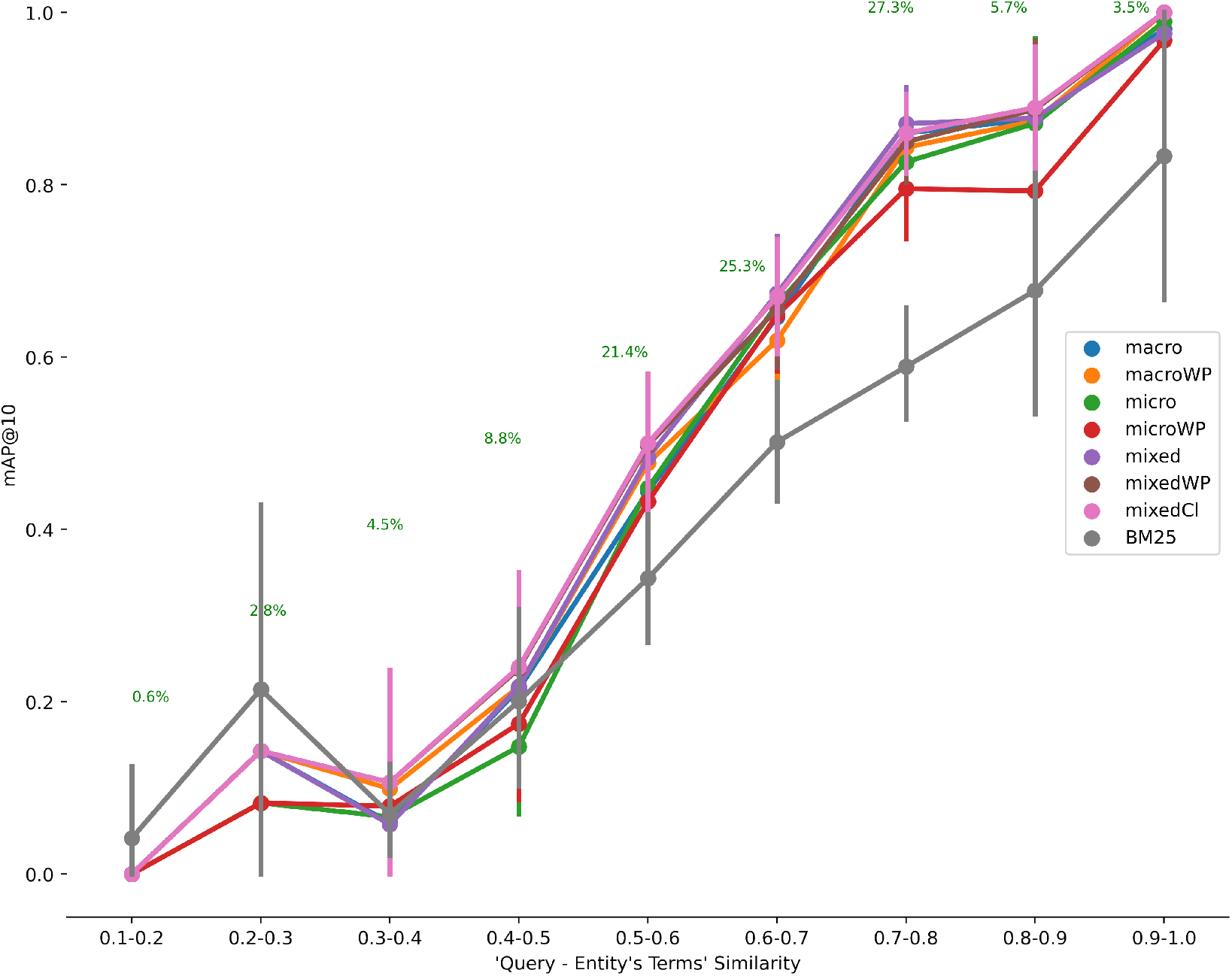
The relationship between the similarity of terms in the query with those in the entity to mAP@10 for PMR-CA. The number of entities is 4,652 and the number of test data is 509. Generally CASBERT is better than BM25 when the similarity value is 0.3 and above.

### 4.2 Recommendations and Future Works

#### 4.2.1 Other Domain Implementation

We have shown CASBERT can well represent composite annotated entities as embeddings. Although the data we use come from a repository of biosimulation models, implementation in other domains such as chemistry, pharmacy or medicine is possible as long as they are annotated using RDF and have ontology dictionaries. The entity retrieval process is started by modifying the input query to embedding using a mixed-method or macro method for more straightforward implementation and then measuring query-entity similarity using cosine similarity. Moreover, most of the BERT models we used are pre-trained models without further fine tuning, except query classification modes; therefore, the implementation in other domains with no training data is still accessible.

#### 4.2.2 Search Engine Implementation

The embeddings representing entities now can be managed in a list-like structure. The list-like structure is more straightforward than the standard indexing technique in search retrieval systems such as the inverted index. New embeddings can be easily attached to the list; even deletion, insertion, and replacement require only a minimum effort; therefore, overall maintenance will be cheaper. More importantly, we can avoid traditional search engine complexities, including preprocessing (stemming, case folding, stop word removal, spelling corrections, and lemmatisation), synonyms and abbreviations handlings.

#### 4.2.3 Cross Repositories Search

We estimate that it is possible to find similar information from different repositories with the same domain. Table 3 shows the example of two queries with their results from the PMR and BioModels database. The query ‘concentration of triose phosphate in astrocyte’, ‘triose phosphate’ is correctly mapped to CHEBI:17138 in entities from both repositories, whereas ‘concentration’ and ‘astrocyte’ are mapped to different entities but with interrelated properties. Furthermore, the query ‘ammonium in cytoplasm’ also gives similar results with the mapping of ‘ammonium’ in entities from both queries was CHEBI:28938 ‘, while ‘cytoplasm’ as FMA:66836 (Portion of cytosol) in the PMR and GO:0005737 (cytoplasm) at BioModels database. These results suggest that multiple repositories can be combined in a single search system to complement each other and be used for possible confirmation and reuse of models across repositories.

**Table 3.** The example of entities retrieved from different repositories, the PMR and BioModels database. Those entities have similarities in the ontology classes related to the queries.

| Query | Result |  |
| --- | --- | --- |
|  | PMR | BioModels database |
| concentration of triose phosphate in astrocyte | **cloutier_2009.cellml#GAPg_GAPg**<br>FMA:54537 Astrocyte<br>OPB:00340 Concentration of chemical<br>CHEBI:17138 glyceraldehyde 3-phosphate | **BIOMD0000000565.rdf#metaid_40**<br>OPB:00592 Chemical molar flow rate<br>CHEBI:32816 pyruvic acid<br>CHEBI:17138 glyceraldehyde 3-phosphate<br>GO:0005829 cytosol |
| ammonium in cytoplasm | **weinstein_1995.cellml#Concentrations.C_int_NH4**<br>OPB:00340 Concentration of chemical<br>CHEBI:28938 ammonium<br>FMA:66836 Portion of cytosol | **BIOMD0000000470.rdf#metaid_1155**<br>OPB:00592 Chemical molar flow rate<br>CHEBI:28938 ammonium<br>GO:0005576 extracellular region<br>GO:0005737 cytoplasm |

#### 4.2.4 CASBERT for SPARQL

SPARQL and CASBERT have a similar intent to get information from RDF documents. SPARQL is rigid and can extract precise details as long as the information about the ontology and structure of the RDF document is known. In comparison, CASBERT is relaxed in exploring information freely where ontology knowledge is not mandatory. Therefore, both methods are not interchangeable, but CASBERT can supplement SPARQL to understand the structure of the RDF document and the ontology classes involved.

#### 4.2.5 Future Works

Henceforth, we will investigate the application of adaptive *w*_*p*_ to calculate the participation of predicates to entity embedding based on the query, as we suggest that this can improve performance. Further study in cross-repositories retrieval also needs to be considered; hence, it can promote reusability between repositories. Finally, the availability of a search engine and API to explore biosimulation models will be beneficial.

## 5 CONCLUSION

The increasing availability of composite annotation to describe entities in biosimulation models requires a simple tool to access information inside by ordinary users. We propose CASBERT, a BERT based method providing keyword-based searching that effectively manages composite annotations and retrieves entities using a text query. This effectiveness is achieved by converting the entities’ composite annotations to embeddings and organising them in a list-like structure; therefore, adding, deleting, inserting, and modifying embedding is cheaper. Getting relevant entities using a previously converted query to an embedding is straightforward with this structure. Using query-entities pairs test data extracted from the PMR and BioModels database, empirically, CASBERT can retrieve better than bag-of-words methods such as BM25. It can potentially give a better user experience than the traditional approach. In the future, we are interested in developing a cross-repositories search engine to encourage biosimulation model reuse between different repositories.

## ACKNOWLEDGMENTS

The authors acknowledge financial support by the Aotearoa Foundation, Auckland Bioengineering Institute, and the National Institutes of Health [grant P41 GM109824].

## DATA AVAILABILITY STATEMENT

The source code, experiment setups and dataset generated for this study can be found in https://github.com/napakalas/casbert

## Footnotes

1 https://www.dbpedia.org/

2 https://yago-knowledge.org

3 https://www.wikidata.org/

4 https://github.com/napakalas/casbert

5 https://models.physiomeproject.org/workspace/5af

6 https://github.com/makcedward/nlpaug

