## Supplemental Tables and Figures for "CASBERT: BERT-Based Retrieval for Compositely Annotated Biosimulation Model Entities"

### Supplementary Material

This document is supplementary material for CASBERT: Composite Annotation Search Based on BERT. Source code, dataset, and experimental setup are available at https://github.com/napakalas/casbert.

#### 1 BIOSIMULATION MODEL - COMPOSITE ANNOTATION QUERY (BM-CAQ)

**Table S1.** Example of query-variables data in the query set in the biosimulation models in the PMR. This set contains a list of queries, the variables associated with the query, and the highest similarity values between the query and the variable variables.

| Query | Variable ID | Max Score |
|---|---|---|
| Calcium-activated potassium current | ’VarId-26893’, ’VarId-19743’ | 0.733406484 |
| Concentration of sodium in the part of cytosol | ’VarId-19766’, ’VarId-26916’ | 0.804430664 |
| Calcium concentration in bulk cytosol | ’VarId-112787’, ’VarId-112086’, ’VarId-64043’ | 0.726958036 |
| Volume of the cytosolic fluid in the diadic space | ’VarId-112102’, ’VarId-112104’, ’VarId-112782’ | 0.715647697 |
| Total cytosolic volume | ’VarId-117177’, ’VarId-112102’, ’VarId-112104’, ’VarId-112782’ | 0.723076522 |

**Table S2:** An example of variables in the PMR annotated with an ontology class with its predicates.

| Variable ID | Class ID | Class Name | Predicate |
|---|---|---|---|
| VarId-26893 | GO:0015269 | calcium-activated potassium channel activity | ’isComputationalComponentFor’, ’physicalPropertyOf’, ’hasMediatorParticipant’, ’hasPhysicalEntityReference’, ’hasPhysicalDefinition’ |
| | OPB:00318 | Charge flow rate | ’isComputationalComponentFor’, ’hasPhysicalDefinition’ |
| VarId-19743 | GO:0015269 | calcium-activated potassium channel activity | ’isPropertyOf’, ’hasMediatorParticipant’, ’hasPhysicalEntityReference’ |
| | OPB:00318 | Charge flow rate | ’isVersionOf’ |
| VarId-19766 | CHEBI:29101 | sodium(1+) | ’isPropertyOf’ |
| | FMA:226054 | Cytosol of neuron | ’isPropertyOf’, ’isPartOf’ |
| | OPB:00340 | Concentration of chemical | ’isVersionOf’ |
| VarId-26916 | CHEBI:29101 | sodium(1+) | ’isComputationalComponentFor’, ’physicalPropertyOf’, ’hasPhysicalDefinition’ |
| | FMA:226054 | Cytosol of neuron | ’isComputationalComponentFor’, ’physicalPropertyOf’, ’part_of’, ’hasPhysicalDefinition’ |
| | OPB:00340 | Concentration of chemical | ’isComputationalComponentFor’, ’hasPhysicalDefinition’ |
| VarId-112782 | FMA:66836 | Portion of cytosol | ’isPropertyOf’, ’is’ |
| | FMA:14067 | Cardiac myocyte | ’isPropertyOf’, ’isPartOf’, ’is’ |
| | OPB:00154 | Fluid volume | ’isVersionOf’ |
| VarId-112086 | OPB:00340 | Concentration of chemical | ’isVersionOf’ |
| | FMA:66836 | Portion of cytosol | ’isPropertyOf’, ’isPartOf’ |
| | FMA:14067 | Cardiac myocyte | ’isPropertyOf’, ’isPartOf’, ’isPartOf’, ’is’ |

*Continued on next page*

Table S2 – *Continued from previous page*

| Variable ID | Class ID | Class Name | Predicate |
|---|---|---|---|
| | CHEBI:29108 | calcium(2+) | ’isPropertyOf’ |
| VarId-64043 | OPB:00340 | Concentration of chemical | ’isVersionOf’ |
| | FMA:66836 | Portion of cytosol | ’isPropertyOf’, ’isPartOf’ |
| | FMA:14067 | Cardiac myocyte | ’isPropertyOf’, ’isPartOf’, ’isPartOf’, ’is’ |
| | CHEBI:29108 | calcium(2+) | ’isPropertyOf’, ’is’ |
| VarId-112102 | FMA:66836 | Portion of cytosol | ’isPropertyOf’, ’is’ |
| | FMA:14067 | Cardiac myocyte | ’isPropertyOf’, ’isPartOf’, ’is’ |
| | OPB:00154 | Fluid volume | ’isVersionOf’ |
| VarId-112104 | FMA:14067 | Cardiac myocyte | ’isPropertyOf’, ’isPartOf’, ’isPartOf’, ’is’ |
| | FMA:66836 | Portion of cytosol | ’isPropertyOf’ |
| | OPB:00154 | Fluid volume | ’isVersionOf’ |
| VarId-112787 | OPB:00340 | Concentration of chemical | ’isVersionOf’ |
| | FMA:66836 | Portion of cytosol | ’isPropertyOf’, ’isPartOf’, ’is’ |
| | CHEBI:29108 | calcium(2+) | ’isPropertyOf’ |
| | FMA:14067 | Cardiac myocyte | ’isPropertyOf’, ’isPartOf’, ’isPartOf’, ’is’ |
| VarId-117177 | OPB:00154 | Fluid volume | ’isVersionOf’ |
| | FMA:66836 | Portion of cytosol | ’isPropertyOf’ |

**Table S3.** Example of query-variables data in the query set in the biosimulation models in the BioModels database. This set contains a list of queries, the variables associated with the query, and the highest similarity values between the query and the variable variables.

| Query | Variable ID | Max Score |
|---|---|---|
| Extracellular glucose kinetics | 'BIOMD0000000051.rdf#metaid_68', 'BIOMD0000000061.rdf#metaid_27', 'BIOMD0000000565.rdf#metaid_75' | 0.720707416 |
| squalene epoxidase (NADP) | 'BIOMD0000000471.rdf#metaid_908', 'BIOMD0000000473.rdf#metaid_972', 'BIOMD0000000472.rdf#metaid_909', 'BIOMD0000000472.rdf#metaid_908', 'BIOMD0000000496.rdf#metaid_4246', 'BIOMD0000000497.rdf#metaid_4262', 'BIOMD0000000471.rdf#metaid_909' | 0.729963005 |
| Factor Xa lipid binding | 'BIOMD0000000332.rdf#metaid_90', 'BIOMD0000000334.rdf#metaid_86', 'BIOMD0000000333.rdf#metaid_62' | 0.709653497 |

**Table S4:** An example of variables in BioModels database annotated with an ontology class with its predicates.

| Variable ID | Class ID | Class Name | Predicate |
|---|---|---|---|
| BIOMD0000000051.rdf#metaid_68 | OPB_00592 | Chemical amount flow rate | 'isVersionOf' |
| | CHEBI:4167 | D-glucopyranose | 'isPropertyOf', 'hasSinkParticipant', 'hasPhysicalEntityReference' |
| | GO:0005576 | extracellular region | 'isPropertyOf', 'hasSinkParticipant', 'hasPhysicalEntityReference', 'isPartOf' |
| BIOMD0000000061.rdf#metaid_27 | OPB_00592 | Chemical amount flow rate | 'isVersionOf' |
| | GO:0015758 | glucose transmembrane transport | 'isPropertyOf', 'isVersionOf' |
| | CHEBI:17234 | glucose | 'isPropertyOf', 'hasSourceParticipant', 'hasPhysicalEntityReference' |
| | GO:0005576 | extracellular region | 'isPropertyOf', 'hasSourceParticipant', 'hasPhysicalEntityReference', 'isPartOf', 'isVersionOf' |
| | GO:0005829 | cytosol | 'isPropertyOf', 'hasSourceParticipant', 'hasPhysicalEntityReference', 'isPartOf', 'isVersionOf' |
| BIOMD0000000565.rdf#metaid_75 | OPB_00592 | Chemical amount flow rate | 'isVersionOf' |
| | CHEBI:17234 | glucose | 'isPropertyOf', 'hasSinkParticipant', 'hasPhysicalEntityReference', 'isVersionOf' |
| | GO:0005576 | extracellular region | 'isPropertyOf', 'hasSinkParticipant', 'hasPhysicalEntityReference', 'isPartOf', 'isVersionOf' |
| BIOMD0000000471.rdf#metaid_908 | OPB_00592 | Chemical amount flow rate | 'isVersionOf' |
| | CHEBI:57945 | NADH(2-) | 'isPropertyOf', 'hasSourceParticipant', 'hasPhysicalEntityReference' |
| | CHEBI:15441 | (S)-2,3-epoxysqualene | 'isPropertyOf', 'hasSinkParticipant', 'hasPhysicalEntityReference' |

*Continued on next page*

Table S4 – *Continued from previous page*

| Variable ID | Class ID | Class Name | Predicate |
|---|---|---|---|
| | P32476 | Squalene monooxygenase | 'isPropertyOf', 'hasMediatorParticipant', 'hasPhysicalEntityReference' |
| | CHEBI:57540 | NAD(1-) | 'isPropertyOf', 'hasMediatorParticipant', 'hasPhysicalEntityReference' |
| | CHEBI:15379 | dioxygen | 'isPropertyOf', 'hasSourceParticipant', 'hasPhysicalEntityReference' |
| | CHEBI:15440 | squalene | 'isPropertyOf', 'hasSourceParticipant', 'hasPhysicalEntityReference' |
| | GO:0005737 | cytoplasm | 'isPropertyOf', 'hasSourceParticipant', 'hasPhysicalEntityReference', 'isPartOf' |
| | GO:0005576 | extracellular region | 'isPropertyOf', 'hasSourceParticipant', 'hasPhysicalEntityReference', 'isPartOf' |
| BIOMD0000000473.rdf#metaid_972 | OPB_00592 | Chemical amount flow rate | 'isVersionOf' |
| | CHEBI:15441 | (S)-2,3-epoxysqualene | 'isPropertyOf', 'hasMediatorParticipant', 'hasPhysicalEntityReference' |
| | CHEBI:15379 | dioxygen | 'isPropertyOf', 'hasMediatorParticipant', 'hasPhysicalEntityReference' |
| | CHEBI:15440 | squalene | 'isPropertyOf', 'hasSourceParticipant', 'hasPhysicalEntityReference' |
| | CHEBI:57945 | NADH(2-) | 'isPropertyOf', 'hasSourceParticipant', 'hasPhysicalEntityReference' |
| | CHEBI:57540 | NAD(1-) | 'isPropertyOf', 'hasMediatorParticipant', 'hasPhysicalEntityReference' |
| | P32476 | Squalene monooxygenase | 'isPropertyOf', 'hasMediatorParticipant', 'hasPhysicalEntityReference' |
| | GO:0005737 | cytoplasm | 'isPropertyOf', 'hasMediatorParticipant', 'hasPhysicalEntityReference', 'isPartOf' |
| | GO:0005576 | extracellular region | 'isPropertyOf', 'hasMediatorParticipant', 'hasPhysicalEntityReference', 'isPartOf' |

#### 2 QUERY CLASSIFIER TRAINING

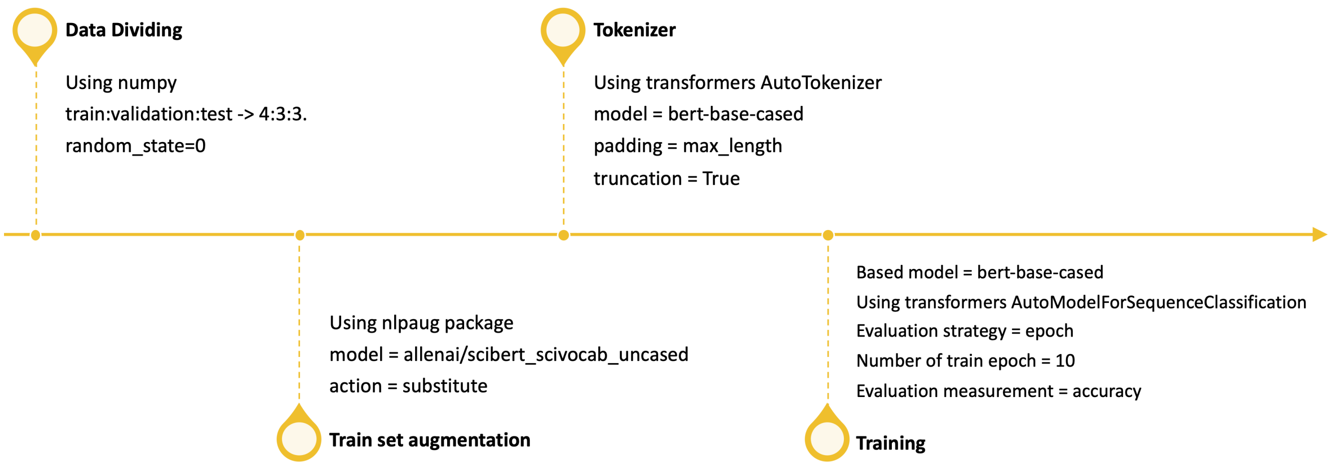


**Figure S1:** Setup for training Query Classifier (QC) model. At first, the dataset is divided into the train, validation, and test sets. The train set is augmented with nlpaug due to the limited number and then tokenised. The number of tokens per query is adjusted to the maximum of tokens used by the BERT model. The training was conducted with ten epoch strategies and evaluated based on accuracy.

#### 3 CASBERT PERFORMANCE FOR BIOMODELS-CA

**Table S5.** CASBERT performance over Biomodels-CA and seven searching strategies compared to the bag-of-words method (BM25) measured using $mAP$@10 and $mRR$. The entities used are only 1,538, which are in BioModels-CA.

| Method | ***noPredicate*** | | ***withPredicate*** | | **combine** | |
|---|---|---|---|---|---|---|
| | **mAP@10** | **mRR** | **mAP@10** | **mRR** | **mAP@10** | **mRR** |
| **BioModels-CA** | | | | | | |
| macro | 0.664723 | 0.654946 | 0.601524 | 0.601969 | 0.663477 | 0.659024 |
| macroWP | 0.625817 | 0.617877 | 0.594065 | 0.600354 | 0.652178 | 0.651609 |
| micro | 0.609719 | 0.607580 | 0.573874 | 0.575732 | 0.616996 | 0.614432 |
| microWP | 0.600927 | 0.600581 | 0.528423 | 0.531402 | 0.591117 | 0.589172 |
| mixed | 0.667825 | 0.659259 | 0.618766 | 0.621912 | 0.675504 | 0.673826 |
| mixedWP | 0.634301 | 0.628718 | 0.609025 | 0.618017 | 0.664714 | 0.665537 |
| mixedCl | **0.669317** | 0.662252 | **0.630671** | **0.636152** | **0.678711** | **0.678250** |
| BM25 | 0.545968 | 0.537574 | 0.535652 | 0.541396 | 0.563251 | 0.561383 |

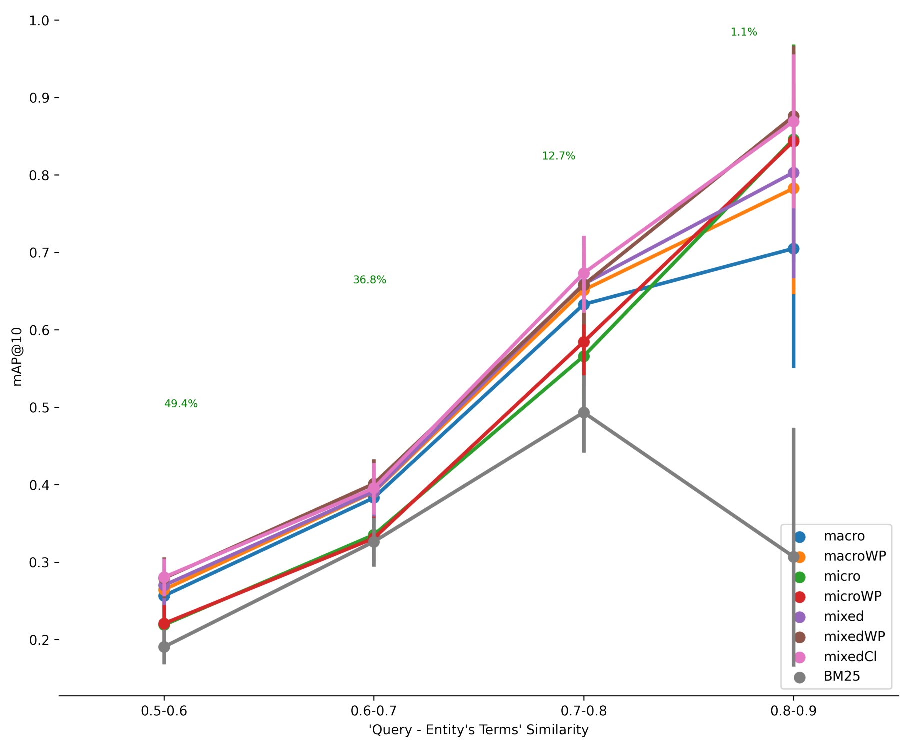


A

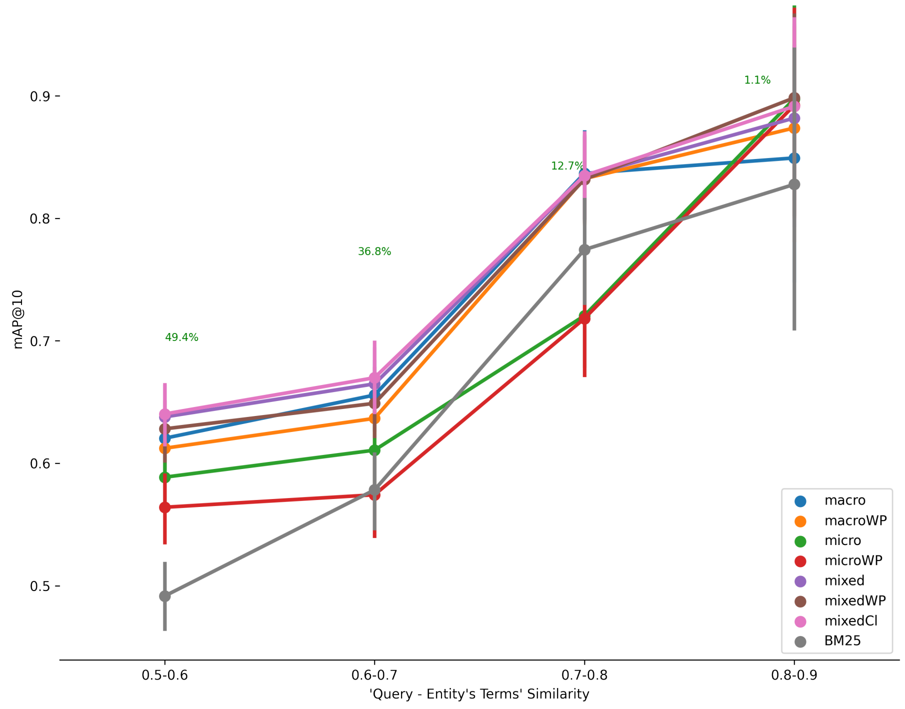


B

**Figure S2:** The relationship between the similarity of terms in the query with those in the entity to mAP@10 for BioModels-CA dataset. The number of test data is 1,777 and (A) compared to all 54,456 entities and (B) compared to entities 1,538 entities appeared in BioModels-CA
